# Population model of Temnothorax albipennis as a distributed dynamical system: I. self-consistent threshold is an emergent property in combination of quorum sensing and chemical perception of limited resource

**DOI:** 10.1101/2021.07.14.452298

**Authors:** Siwei Qiu

## Abstract

House hunting of ant, such as Temnothorax albipennis, has been shown to be a distributed dynamical system. Such a system includes agent-based algorithm [1], with agents in different roles including nest exploration, nest assessment, quorum sensing, and brood item transportation. Such an algorithm, if used properly, can be applied on artificial intelligent system, like robotic swarms. Despite of its complexity, we are focusing on the quorum sensing mechanism, which is also observed in bacteria model. In bacterial model, multiple biochemical networks co-exist within each cell, including binding of autoinducer and cognate receptors, and phosphorylation-dephosphorylation cycle. In ant hunting, we also have ant commitment to the nest, mimicking binding between autoinducer and cognate receptors. We also have assessment ant specific to one nest and information exchange between two assessment ants corresponding to different nests, which is similar process to the phosphorylation-dephosphorylation cycle in bacteria quorum sensing network. Due to the similarity between the two models, we borrow the idea from bacteria quorum sensing to clarify the definition of quorum threshold through biological plausible mechanism related to limited resource model. We further made use of the contraction analysis to explore the trade-off between decision split and decision consensus within ant population. Our work provides new generation model for understanding how ant adapt to the changing environment during quorum sensing.

## I. Introduction

Quorum sensing [2–4] is important systematic stimulus and response that is sensitive to population density of interest. Functionally, ant is using quorum sensing to coordinate behaviors including foraging [5], house hunting [6, 7], and cooperative transport [8]. Other biological system, like bacteria [9], also have similar behavior, like bioluminescence [10], biofilm information [11], virulence [12], and antibiotic resistance [13], which are relying on local density of bacterial population. Similar behavior also seen in group of fish [14], group of bird [15], etc. Similar performances are seen in other distributed system, like those related to vote in political event [16], where competition mechanism is involved and bias the decision. These quorum sensing networks are resembling cognitive function of a brain [17], which is particularly a decision-making process [18, 19] that require collective response to external stimulus and make a decision that is beneficial to the goal of the specific task [20].

In the case of ant, quorum sensing is just part of the story, where other activities, including exploration to search nest, assessment of the found nest, and moving brood item after the decision is made. In the first stage, the searching ant is searching for potential nest due to fact that the nest they had before was destroyed by the environment (raining for example). Once the searching ant found the candidate nest, they go back to the original nest, and pass the information to the other ant through pheromone (as explained in Fig 1), resulting them to go do assessment of this candidate nest. Each assessment ant needs to decide on whether they commit to this nest or not based on their comparison of the quality of this candidate nest. Once they decide to commit, they start to recruit more ant to that nest through pheromone [21]. The quorum sensing starts here, because it is observed that once the quorum sensing criterion is met, meaning that the proportion of the ant commit to this nest meat the threshold, the recruit ant stops recruiting more ant, and start to transport the brood item to that nest, and decision already made. This process is a positive feedback, which doomed to happen for single nest model, but a bit more complicated in multiple nest case. In the case of two nests for example, due to competition mechanism, there are negative feedback in each of the nest due to some process that the ants communicate and exchange information, leading to the shift of commitment. The competition will lead to either winner take all [22] or “synchronization” [23, 24]. The scheme of such kind of experiment is illustrated in Fig 2. Of course, the winner-take-all case is mostly seen when the two nest-candidates are with different quality, but synchronization is seen for case where two nests candidate are with same quality.

**Fig 1.**
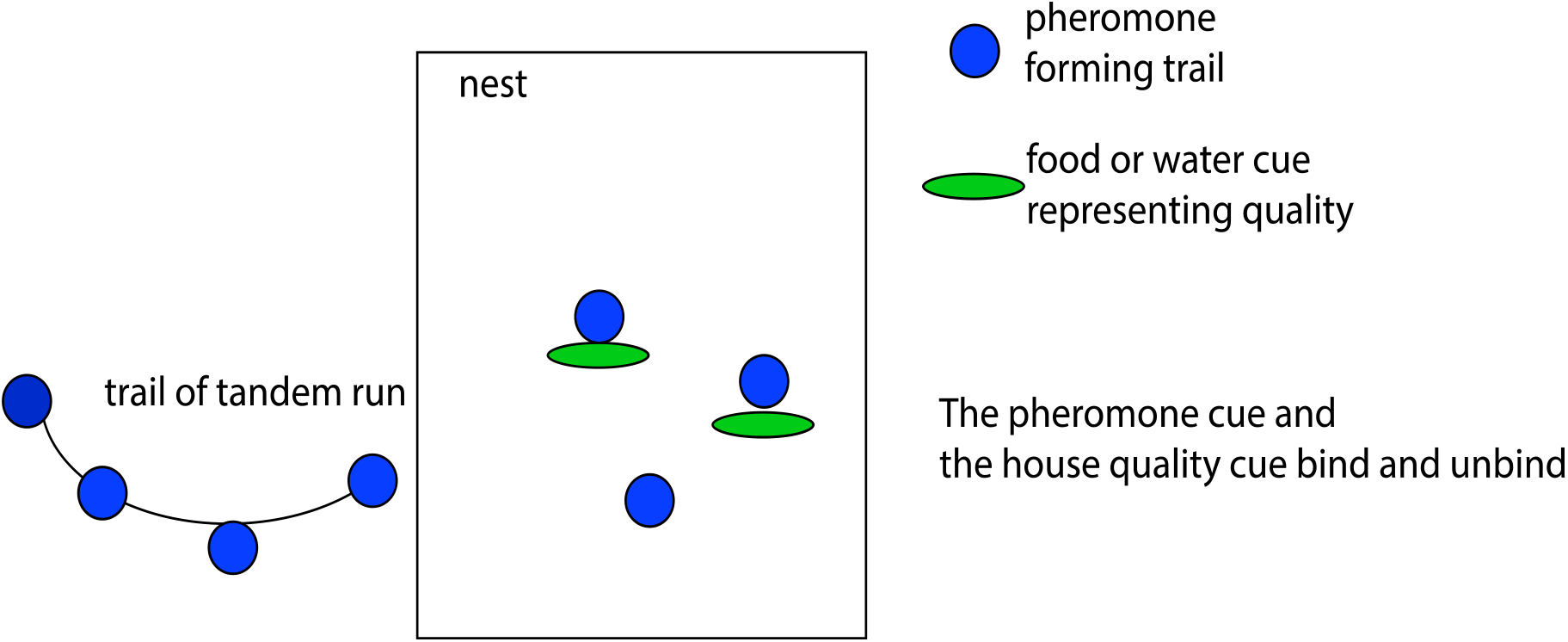
The recruit ant releases the pheromone cues to form a trail outside nest so that follower ant can find the nest. Within the nest, there is binding and unbinding of the pheromone of ant with the house quality cue (can be food and water). The ant house hunting is promoted by these chemical stimuli during their navigations [25]. Just like the released vesicle to bind the neural transmitter in the neuroscience literature, the binding site of house quality cues are finite number. With this constrain of “limited resource”, we have a biological plausible mechanism for threshold. This gives a theoretical framework for the “plasticity” rule of house hunting ant.

**Fig 2.**
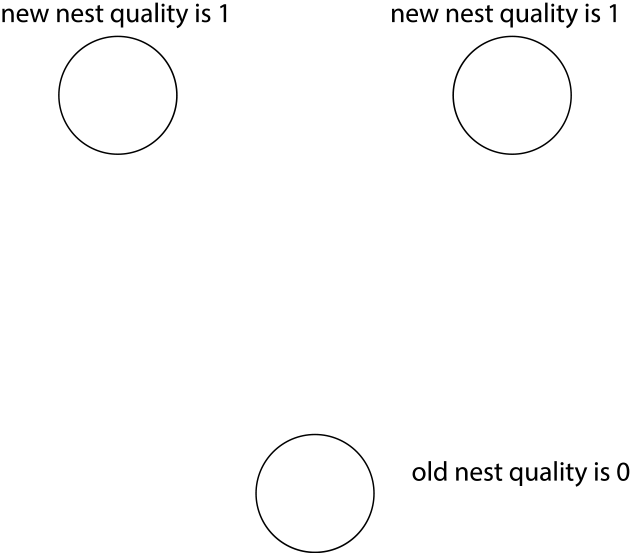
Typical experiment [26] set up for house hunting ant with two new nests. The old nest is destroyed, and the ants are forced to explore and find a new house. The phenomenon that the ant invests long time to make decision is supported by global converging theory with contraction.

Majority of the ant quorum sensing are based on deterministic ordinary differential equations (ODEs) introduced by SC Prat [3]. Recent effort also been put into spatial models. From dynamical distributed system’s point of view, we are mainly considering two specific feedbacks, the positive feedback from recruit of ant that commit to same nests as their lead, and the negative feedback coming from parallel pathways, including reject by chance to commit to same nest as their lead, and the “rumor” transferred from searching ant who has information of other “better” nests. Two possible resulting dynamics can be observed, one is switching dynamics, and another is oscillation. In this paper, we are focusing on explaining switching dynamics, which is the adaptation with the changing environment. On one hand, we show that the network is robust under “rumor” passing through the network, on the other hand, we show that the network is flexible and can adapt to the correct decision even when the environment (here we focus on quality of the nests candidates) are changing.

Recent experiment studies the quorum sensing in the ant house hunting, with mediocre and high-quality nests, and observed speed-accuracy trade-off. However, the theory behind this phenomenon is missing. In this work, we make use of dynamical system and contract analysis, and the relation between global converge and the parameters robustly. As a side product, we show that the threshold is not independent parameter like most of the literature illustrated, but it is related to other parameters, like state transition rate, through self-consistent equation.

The structure of the paper is as follows. In Sec. II, we derive the condition for the global convergence of the ant house hunting system and give prediction on the converging rate through eigenvalue of the system. In sec. III, we discuss further on the relation between threshold of each nest and the parameters.

This leads to discussion on how changing threshold will emerge due to environment change related to the house quality.

## II Global convergence

Derive the contraction theory [2, 27, 28]. In general, we are considering *n* dimension dynamical system:

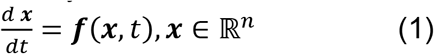

Here, map *f* : ℝ^*n*^ → ℝ^*n*^ is smooth nonlinear vector field, we then can define *L^p^* norms for vector and matrix *x* and A. Also, can define matrix measure *μ* as:

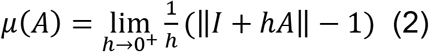

One can define *L^p^* norms of *μ*, and the following theorem hold:

### Theorem 1.

The n-dimensional dynamical system is said to be contracting if any two trajectories, starting from different initial conditions, converge exponentially to each other. A sufficient condition for a system to be contracting is the existence of some matrix measure *μ* for which there exists a constant *λ* > 0 such that:

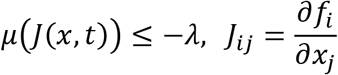

For all **x**, t. The scalar *λ* defines the rate of contraction.

In physics, this the contraction is recognized as entropy increasing, which is a requirement of the entropy increasing theorem. We are interested in this because in experiment we already see surprising result that when two equal quality nests are presented, the decision is slower. In this sense, the contraction rate is minimized when there exists symmetry in the system.

The contraction theory gives us a unified description of the system with just single nest dynamics, when the global converge happen. In next section, we will first explore the single nest dynamics, where we will find the definition of important variable, like threshold of quorum sensing.

## III Quorum Threshold

### 1. Single nest model

We consider the following simple process for single nest model:

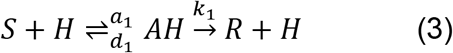

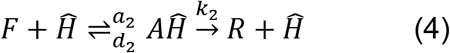

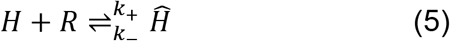

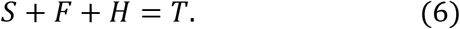

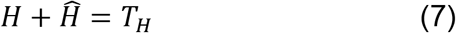

Here, *S* represent the searching ant, *F* represent the follower that follow leading ant (the recruit ant) in tandem run, *H* represent the site in the nest which is not explored yet (or less concentration of the pheromone stimuli), and 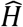 represent the site on nest where recruiter left some pheromone cue. We change variable, and denote the number of different type of ants using *μ* = [*R*], *w* = [*s*], *w*\* = [*R*\*], *r* = [*H*], 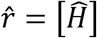 *ν* = [*AH*], 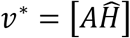. We then construct the following differential equations:

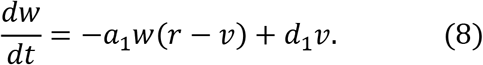

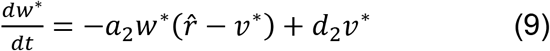

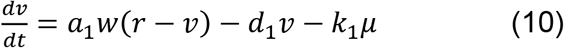

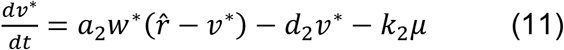

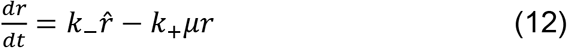

Solving these equations, and treating variable *r* as fast variable, we can get the stationary solution:

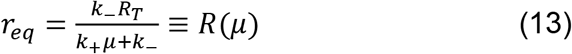

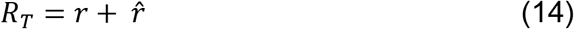

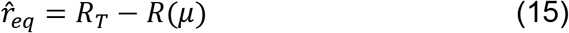

We also treat *ν* and *ν*\* as fast variable, and can solve their equilibrium solution, and get 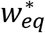 as solution of a quadratic equation:

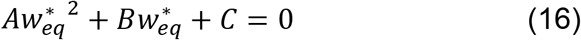

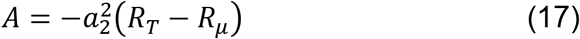

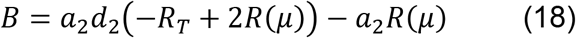

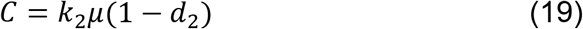

We then get the solution of 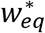 as solution:

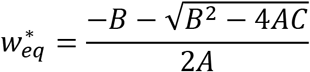

We check with numerical calculation and explore the parameter, and found that under appropriate parameter choice, the recruit ant number is well constrained by boundary condition imposed by the stationary solution of the number of quality cues in the nests. Within such a boundary, the upper bound of the stationary solution of assessment ant is always reasonable and close to zero. We then claim that limited resource of the quality cue within a finite volume nest provide biological mechanism of the threshold of recruit ant number. Decision is made when there is recruiter for more assessment ant, but assessment is not necessary. At this moment, the recruit ant will start transit to do transportation, and keep the total number of recruiters in a reasonable range. They will only start recruit again when “urgent” situation occurs, for example, when 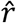 reduce unexpectedly. If this happens, exploration ant will start exploring somewhere else for higher quality nest, meaning with higher 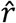.

**Figure 2 A.**
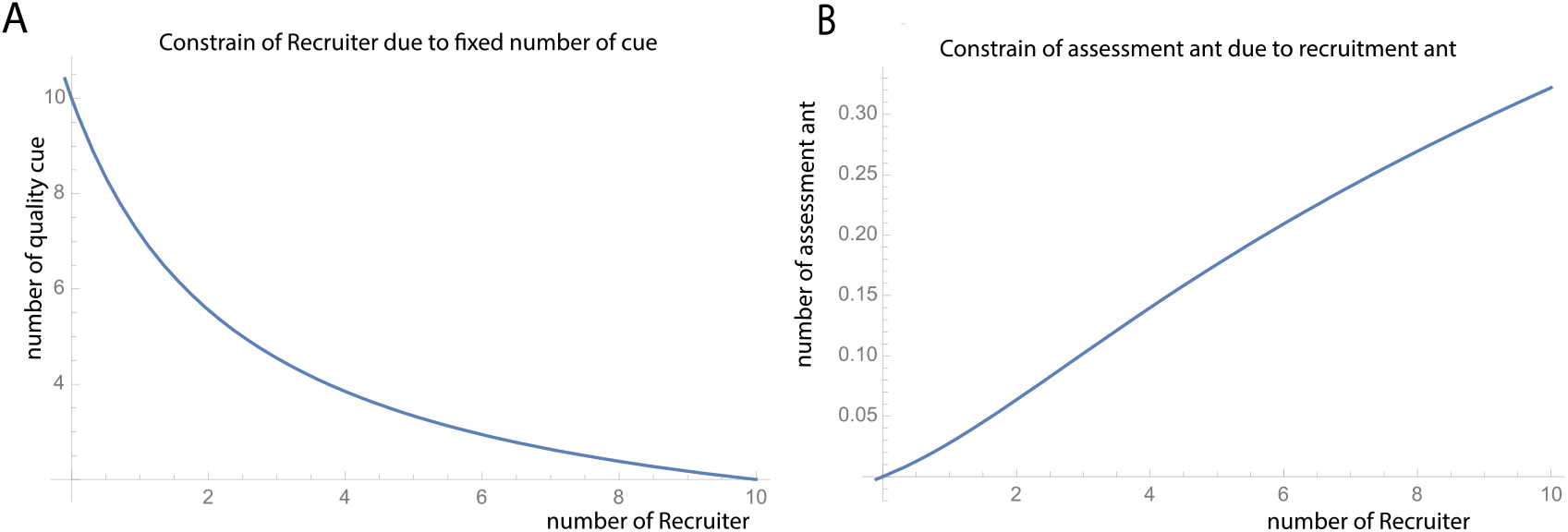
Under appropriate parameter setting, the stationary solution of the number of house quality cue is reducing as the number of recruiter increase. And this provide the requirement that the total number of recruiters having an upper bound, since the number of the quality cue is non-negative. B. The stationary solution of assessment ant (was follower before they reach the nest) is always close to zero and have an upper bound due to upper bound of number of recruiters. In this calculation, we set *R_T_* = 10, since we require *R_T_* ≪ *N_ant_*, where *N_ant_* is total number of ants, usually around 100.

### II Multi-nest model

We argue that a reduced model is needed for ant house hunting model. We borrow the “compound state” idea from chemistry to reduce dynamical variable, since it can represent the recruit ant. We treat the exploring ant as the “common median” of the model, and describe it using synaptic dynamics shared by the whole network (shared memory in distributed algorithm):

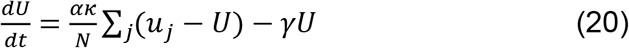

And we define the recruit ant dynamics in the following form:

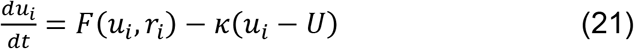

We have dynamics of “binding sites (representing quality of house, related to the pheromone of ants)” in the nests as follows:

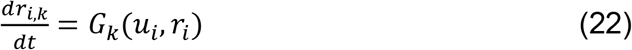

This then reduce the complexity of the model, allowing us to get the main feature of the ant house hunting model.

Following the steps of literature [2], we list the resulting dynamics of reduced model as:

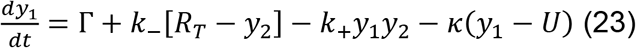

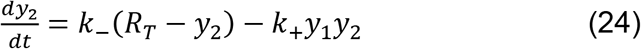

And the Jacobian is:

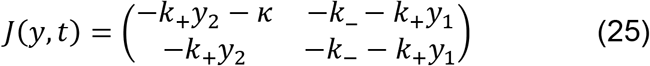

One then can construct the eigenvalue of the symmetric matrix 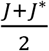, which are *λ*_±_. We then can get the maximum eigenvalue as matrix measure, giving definition of the converge rate. This also explain the minimum rate, meaning slow decision, because this eigenvalue is negative value, so absolute value is small. We therefore can use simple coupled ODE to explore the converged dynamics of the system, and this can predict the converge time, which is related to the eigenvalue of the reduced system.

**Fig 3.**
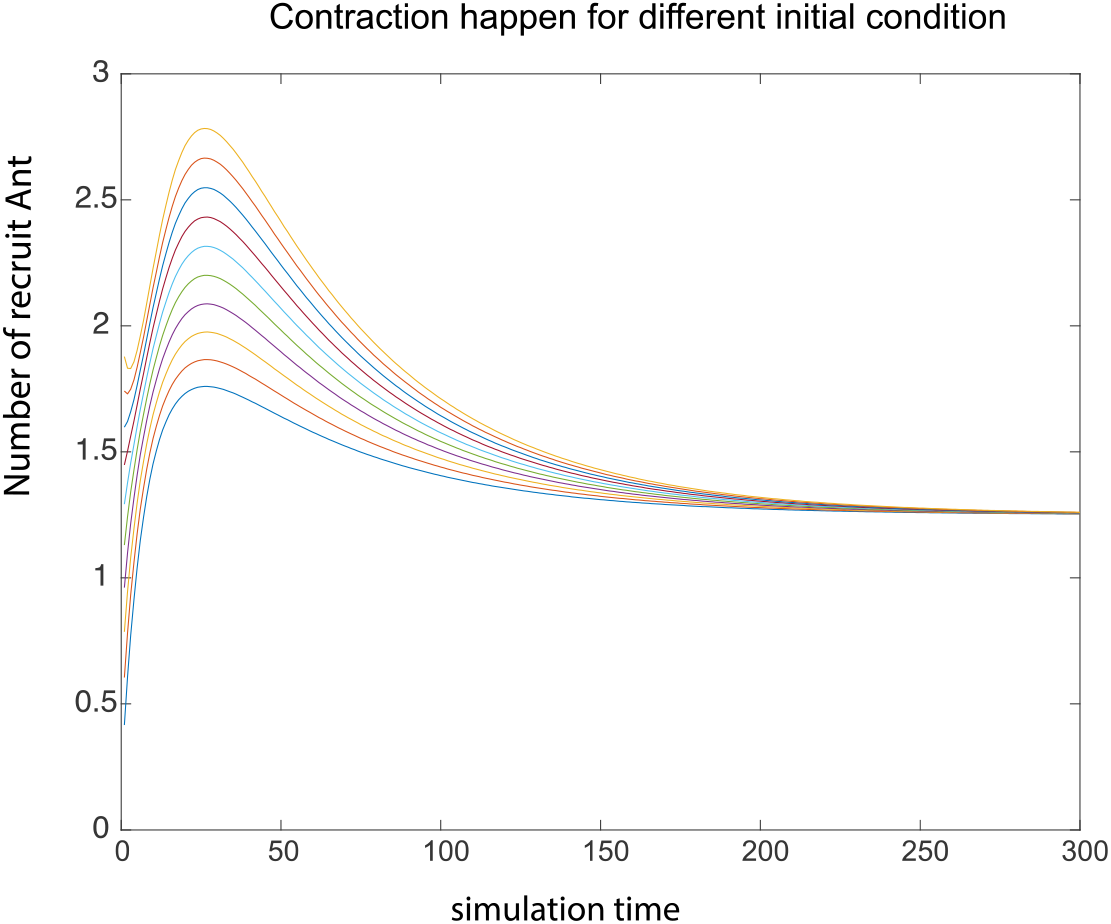
Simulation shows global converge happens, this means we can use mean field theory to explore the property of multi-nest model when there is synchrony, which is like the case of decision in present of two (or more) equal quality nests. We see that although they can have different transient dynamics, the result always converges [27] after enough decision time.

## Discussion

In this paper, we showed that using a reduced model that is repeated explored in bacterial quorum sensing problem, we can get insight of the behavior of the house hunting ant. The global feedback from “common media”, the recruit ant is crucial in decision making. With a slow dynamics of recruit ant, we have dynamics like the synaptic dynamics [29] in neuroscience. In neuroscience, slow dynamics of synaptic modify the communication between neurons, and result in collective decision within network of neurons. In here, the modulation of recruit ant, act as feedback and regulate the dynamics in each nest. The dynamics of the modulation by cues of environment in the nests of ant is similar to neural transmitter dynamics triggered long time scale dynamics of the whole system, which are mimicking the limited resource model in auditory neuroscience [30]. Indeed, the repeated increase of recruiting ant, resulting in a “threshold” of the total recruit ant number, which is the biomarker of the decision being done, which is related to the time constant of the whole system. We revisit some result in bacterial literature, and show that the converge speed, which is related to the time investment in the decision making and the time scale of the whole system, is related to the eigenvalue of the reduced system in context of partial contraction theory. And this explain why contradict to intuition, when two nests are with equal quality, the ant group takes longer time to make decision. Our theory further defines the threshold by solving a self-consistent equation of assessment ant number, which is common technique in the theoretical neuroscience literature [31]. This leads to a new way to build theory of house hunting ant because the previous models are all treating threshold as an independent parameter. We show the correlation between different parameter through computational simulation and showed a more natural in a sense of the limited resource context. In the future, we will further explore the house hunting behavior with present of geometry structure (distance of from the candidate nest to the old nest), which can be same analysis as mean field theory literature for ecology [27, 32] or extension of distributed algorithm literature [33]. This will add in new factor for the decision making and making the merit function more realistic since ant will take the effort into account.

## Acknowledgement

We thank Jiajia Zhao in MIT for providing detailed discussion. We also thank Nancy Lynch in MIT for her class on distributed algorithm and her encouragement on thinking in more theoretical way, which is the reason I start to work on this project. I also thanks university of Pittsburgh for supporting my research.

## Notes

### Competing Interest Statement

The authors have declared no competing interest.

